# A Murine Model of Abductor Insufficiency Accelerates the Development of Hip Osteoarthritis

**DOI:** 10.1101/2022.05.22.492940

**Authors:** Michael B. Geary, Caitlin A. Orner, Helen Shammas, John M. Reuter, Alayna E. Loiselle, Chia-Lung Wu, Brian D. Giordano

## Abstract

Osteoarthritis (OA) of the hip is a common and debilitating painful joint disease. A growing body of evidence suggests that there may be an association between periarticular myotendinous pathology and the development of hip OA. Thus, we hypothesized that a murine model of hip OA could be achieved through selective injury of the abductor complex around the hip. C57BL6/J mice were randomized to sham surgery or abductor injury, in which the myotendinous insertion at the third trochanter and greater trochanter were surgically detached. Mice were allowed free, active movement until sacrifice at either 3 weeks or 20 weeks post-injury. Histologic analyses and immunohistochemical staining (IHC) of the femoral head articular cartilage were performed, along with μCT analysis to assess subchondral bone remodeling. We observed that mice receiving abductor injury exhibited significant OA severity with loss of Type II Collagen staining compared to sham control mice at 20 weeks post-surgery, comparable MMPI13 expression was observed between injury and sham groups. No significant differences in subchondral bone were found on μCT after 20 weeks following injury. Our study suggests a link between abductor dysfunction and the development of hip OA, which are common pathomorphologies encountered in routine orthopaedic clinical practice. Further, this novel animal model may provide a valuable tool for future investigations into the pathogenesis and treatment of hip OA.

## Introduction

Osteoarthritis (OA) affects over 32.5 million people in the United States, and approximately 10% of adults over the age of 45 will suffer from symptomatic hip OA at some point in their life^1^. The hip is the second most commonly affected large joint, after the knee^2^. Hip OA is generally divided into primary idiopathic causes, and secondary causes resulting from an identifiable factor^3^. Secondary degeneration of the hip can result from a number of pathologies, including inflammatory diseases such as rheumatoid arthritis, and has been linked to structural pathomorphologies such as femoral acetabular impingement (FAI)^4^. Despite much study on the causative factors related to the onset and progression of hip OA, the underlying etiology remains to be elucidated.

The hip is a diarthrodial joint that allows for multidirectional range of motion. The periarticular musculature of the hip, in particular the gluteus medius and minimus and their associated tendons, contribute to dynamic stability of the joint. It is well-recognized that altered arthrokinematics negatively influence articular cartilage loading and contributes to the progression of OA. Multiple studies have observed an association between periarticular tendinopathy and hip OA, without definitively establishing causality^5,6^. In a clinical study, a small series of patients with abductor tears were found to demonstrate a high rate of concomitant intra-articular injury^7^. Therefore, these studies provided rationale for considering whether musculotendinous dysfunction or injury surrounding the hip may lead to the development of OA.

Animal models of pathogenesis of hip joint and OA mostly utilize large animals including cannies^8-10^, sheep^11^, and porcine^12^ with the advantage of large animals having similar cartilage morphology, thickness as well as the responses to injury to humans^13-15^. These large animal hip OA models not only have already provided us invaluable insights into hip cartilage physiology and pathophysiology but also serve as essential pre-clinical models for drug development.

Despite these benefits, large animals are not amenable to the power of genetic modification to better define the cellular and molecular etiology of pathogenesis, as can be conducted in mice^15^. Furthermore, there is a paucity of literature on murine hip injury models^16^. The purpose of the present study, therefore, was to establish a novel murine model of hip OA induced by abductor injury and determine whether an association exists between abductor insufficiency and hip OA onset and progression. We hypothesize that abductor injury accelerates the development of hip OA. OA severity, subchondral bone remodeling, as well as anabolic and catabolic biomarkers were evaluated by histology, immunohistochemistry (IHC) and uCT analyses in the current study.

## Materials and Methods

### Animal Model of Abductor Instability

All animal procedures were approved by the University Committee on Animal Research (UCAR) at the University of Rochester. Female C57BL/6J mice (8-10 weeks-old; #00664, Jackson Laboratories, Bar Harbor, ME) weighing on average between 20 and 25 g were anesthetized by intraperitoneal injection of ketamine (60 mg/kg) and xylazine (4 mg/kg). The hind region and right leg were shaved and prepared with alcohol and betadine (povidone-iodine). The proximal femur was approached through a 1 cm lateral skin incision placed over the third trochanter, followed by a direct incision through the fascial layer, exposing the underlying third and greater trochanters.

Mice were randomly divided into two groups. One group received abductor injury, while the other group received sham surgery. The effect of abductor injury on hip OA development was investigated at two time points following surgery: 3 weeks and 20 weeks post-surgery. n = 4-5 mice per group (sham or surgery) per time point. The sham group underwent anesthesia along with incisions of the skin and fascia. A Schematic diagram of the right proximal femur shows the locations of the femoral head (FH), greater trochanter (GT), lesser trochanter (LT), third trochanter (3T), and sciatic nerve (SN) (**Fig. 1A**). Coronal μCT reconstruction of the right hip demonstrates the anatomic positions of the FH and GT (**Fig. 1B**). The injury group had release of the muscular attachments to the 3T (**Fig. 1C**) as well as detachment of the large abductor complex (ABD) from the GT (**Fig. 1D**). During each surgery, careful attention was paid to avoid injury to the SN running posterior to the proximal femur. For all mice, the fascial layer was closed with a single 5-0 nylon suture (Ethicon, Inc.), and the skin was closed with either 7 mm stainless steel wound clips (CellPoint Scientific Inc., Gaithersburg, MD) or a series of interrupted 5-0 nylon sutures. For analgesia, mice were given subcutaneous buprenorphine (0.05 mg/kg) at the time of surgery and each subsequent day over the following 72 hours. Following surgery, mice were returned to their cage and allowed free active motion and weight bearing. Animals were monitored daily until sacrifice.

**Figure 1.**
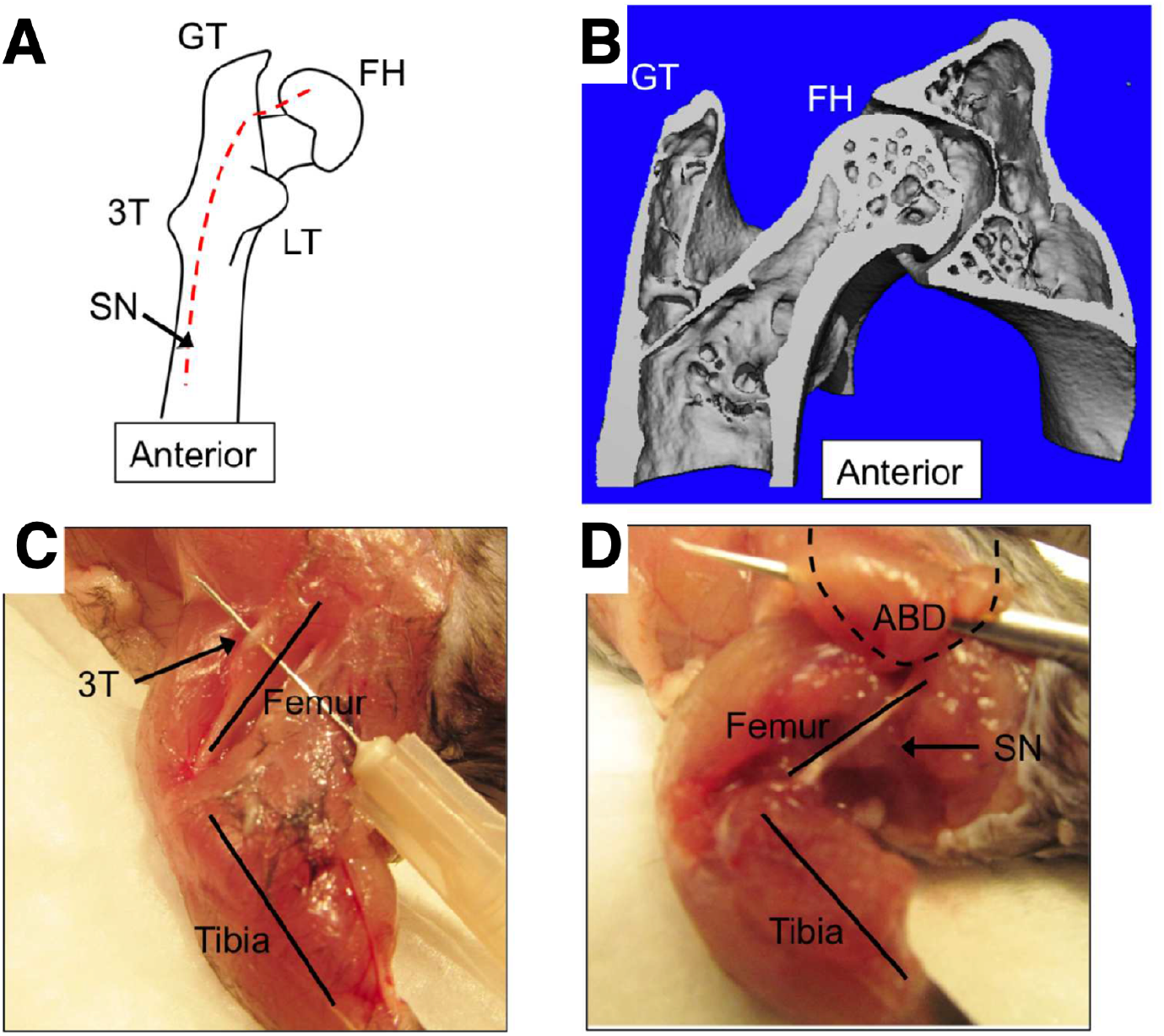
Surgical model of abductor complex injury. (**A**) Schematic of the right proximal femur showing the locations of the femoral head (FH), greater trochanter (GT), lesser trochanter (LT), and third trochanter (3T), as well as the path of the sciatic nerve (SN) running posterior to the femur. (**B**) Coronal reconstruction of the right hip showing the anatomic positions of the femoral head (FH) and greater trochanter (GT). (**C**) Photograph of the left hind limb with skin removed. The femur and tibia are labeled for orientation. The third trochanter (3T) is visible superficially, with a needle placed into the muscular attachments. Attachments running proximal from the third trochanter were removed in this injury model. (**D**) Photograph of the left hind limb highlighting the abductor attachments (ABD) to the greater trochanter (dashed outline). The entire abductor complex was detached in the injury group. The femur, tibia, and sciatic nerve (SN) are labeled for orientation.

### Evaluation of bone tomography

Mice were sacrificed at either 3-weeks or 20-weeks post-surgery (n = 4 -5 per group per time point). Immediately after sacrifice, the hemi-pelvis and femur were prepared by removing the skin and excess soft tissue, leaving the hip and the major muscular attachments around the pelvis and femur. The samples were fixed for 72 hours in 10% neutral buffered formalin (NBF), then washed, placed in 70% EtOH, and subjected to the μCT scanning. Hips were scanned using the VivaCT 40 system (Scanco Medical, Basserdorf, Switzerland) at highest resolution with a pixel size of 10.5 μm to image bone. An integration time of 300 ms and an X-ray voltage of 55 kVP was used. The hip was captured and segmented using a scanning threshold of 320 to identify bone. The femoral head was analyzed for trabecular bone (excluding cortical) at a threshold of 310. Views of the femoral head and acetabulum were analyzed using a threshold of 320. The Gaussian method was used for noise reduction (sigma 0.8 with support value of 1 pixel).

### Evaluation of OA severity

Following 72 hours fixation and μCT scanning, samples were decalcified in 14% (pH 7.2) EDTA at room temperature for 2 weeks. Samples were then dehydrated and embedded in paraffin with the posterior surface of the hip facing down. Using the SN as a landmark, serial sections (thickness = 5 μm) were taken through the preserved hip and surrounding soft tissue in a coronal plane. Three different levels were cut through the joint, with five sections taken at each level.

Safranin O / Fast Green staining was used to visualize changes in the hip articular structure and proteoglycan content of the mice at 3 weeks and 20 weeks post-surgery. Slides were baked overnight at 60°C, then deparaffinized and rehydrated through graded ethanol. After air drying, slides were stained in Weigert’s Hematoxylin for 7 minutes (Equal parts Solution A, CAS# 517-23-2 and Solution B, CAS# 7705-08-0) then rinsed with running tap water. Slides were stained with 0.02% Fast Green (CAS# 2353-45-9) for 3 minutes, rinsed in 1% acetic acid for 10 seconds, then stained with 1% Safranin O (CAS# 477-23-6) for 5 minutes. After a quick rinse in 0.5% acetic acid and rinses with double-distilled water, slides were then air dried and cover slipped.

OA severity was determined by modified Mankin scoring system as previously described^17^. Three independent, blinded graders assessed sections for degenerative changes in the hip articular cartilage. Scores including articular structure (0-11), tidemark (0-3), loss of proteoglycan staining (0-8), and chondrocyte clones (0-2) were averaged among graders for the hip femoral head, resulting in total scores between 0 and 24.

### Immunohistochemistry

Immunohistochemistry (IHC) was performed for Collagen Type II A1 (COL2A1) and Matrix Metalloproteinase 13 (MMP13). For both protocols, slides were baked overnight at 60°C, then deparaffinized and rehydrated through graded alcohols. IHC labeling for MMP13 (Thermo Fischer Scientific, Waltham, MA. Catalog #MS-825P) began with enzymatic antigen retrieval using hyaluronidase (Sigma-Aldrich, St. Louis, MO; H-3506) for 10 minutes in a 37°C water bath. Endogenous peroxidase was then quenched for 30 minutes using endogenous blocking reagent (Dako North America, Carpinteria, CA; S2003). After rinses with deionized water and PBS-T, slides were blocked for 30 minutes with 5% Normal Horse Serum (NHS; Vectastain Elite ABC Kit (Mouse IgG), Vector Laboratories, Burlingame, CA; PK-6102), then with the Blocking Endogenous Antibody Technology (BEAT) kit (Invitrogen Corporation, Camarillo, CA; 50-300). Overnight incubation with the primary antibody (Thermo Fischer Scientific, Waltham, MA; MS-825P; 1:200) was done at 4°C, with control slides incubated with mouse IgG at the same concentration (1ug/mL). On the second day, sections were incubated with the secondary antibody (Vector PK-6102), then ABC reagent (Vector PK-6102). Color was detected using Vector DAB ImmPACT kit (Vector SK-4105), then counterstained with hematoxylin.

IHC labeling for COL2A1 was performed as follows. Enzymatic antigen retrieval was performed for 10 minutes in a 37°C water bath using pepsin (Sigma-Aldrich, P7000). After rinses, sections were blocked with endogenous blocking reagent (Dako, S2003) for 30 minutes, then with 5% NHS (Vector PK-6102) for 30 minutes. The primary antibody (Thermo Scientific, MS235-P; 1:100) was left to incubate at 4°C overnight, with control slides incubated with the same concentration of mouse IgG (2ug/mL). On the second day, sections were incubated with the secondary antibody (Vector PK-6102) and ABC reagent (Vector PK-6102), then color was detected with the Vector DAB ImmPACT kit (Vector SK-4105) counterstained with hematoxylin.

### Statistics

To determine how surgery, time (following surgery), and their interaction (surgery x time) affected hip OA severity, as well as subchondral bone remodeling, data were analyzed by two-way repeated measures ANOVA following by Sidak’s multiple comparisons test (α = 0.05) using GraphPad Prism version 9. Values are expressed as mean ± SD.

### Source of Funding

This study was supported by NIH AR075899 (CLW), OREF Etiology of Hip Osteoarthritis Grant (CLW & BG) as well as Goldstein Award (BG) through the University of Rochester’s Department of Orthopaedics and Rehabilitation. MBG was supported by CTSA (TL1 TR000096) from NIH/NCATS. CAO was supported by the University of Rochester Office for Medical Education Year-Out Research Fellowship. The HBMI and BBMTI cores are supported by NIH/NIAMS P30 AR069655.

## Results

### Mice receiving abductor surgery demonstrated significantly increased OA severity compared to sham group 20 weeks following injury

Safranin-O is a cationic dye that stains proteoglycans, and intensity of red Safranin-O staining is a proxy for proteoglycan content. Loss of proteoglycan content is a clinical hallmark of OA^18^.

Hips from the sham and injured groups were stained at 3 weeks and 20 weeks post-injury with Safranin-O/Fast Green to visualize proteoglycans. After 20 weeks, mice in the injured group demonstrated loss of proteoglycan staining of the articular surface relative to sham, although articular cartilage surface was comparable between two groups (**Fig. 2A-B**). No significant differences in OA severity was observed between sham and injured groups at 3 weeks post-injury. Furthermore, sham mice had similar OA severity at 3 weeks and 20 weeks post-surgery.

**Figure 2.**
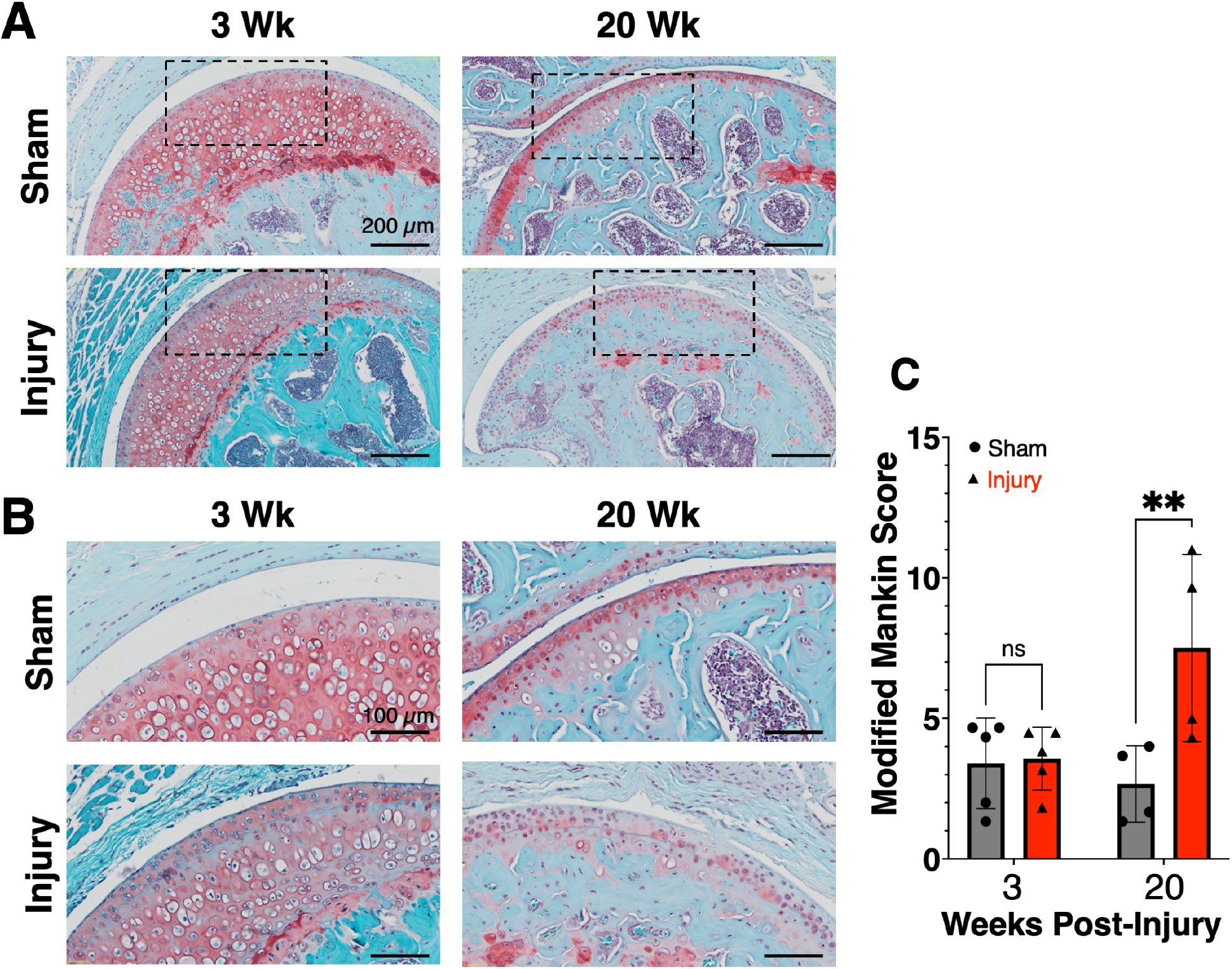
Abductor injury results in loss of proteoglycan staining 20 weeks post-surgery. Coronal sections through the hip were stained with Safranin O/Fast Green, in which proteoglycan stains red. (**A**) Sections are shown for sham and injured groups at 3 and 20 weeks after surgery. Loss of proteoglycan staining is evident in the injured group after 20 weeks relative to sham group. (**B**) Increased magnification of the areas indicated by dashed rectangles in (**A**). (**C**) Modified Mankin scores shows increased OA severity of the mice receiving surgery 20 weeks post-injury as compared to corresponding sham group. Two-way repeated measured ANOVA following by Sidak’s multiple comparisons test. Mean ± SD. ns: No statistically significant was detected.

### Mice receiving abductor surgery exhibited decreased COL2A1 staining but comparable MMP13 staining in the articular cartilage relative to sham mice at 20 weeks following abductor injury

IHC labeling for COL2A1 and MMP13 in articular cartilage of the femoral head was performed at 20 weeks post-injury as this was the timepoint when OA was evident in the mice receiving surgery. In the sham group, a superficial layer of unmineralized cartilage is characterized by pericellular staining of COL2A1 (**Fig. 3A-B**). In the deeper layers, staining intensity is increased and more evenly distributed throughout the extracellular matrix. Hips from the injury group show a notable decrease in staining intensity in the uncalcified articular cartilage, relative to sham mice.

**Figure 3.**
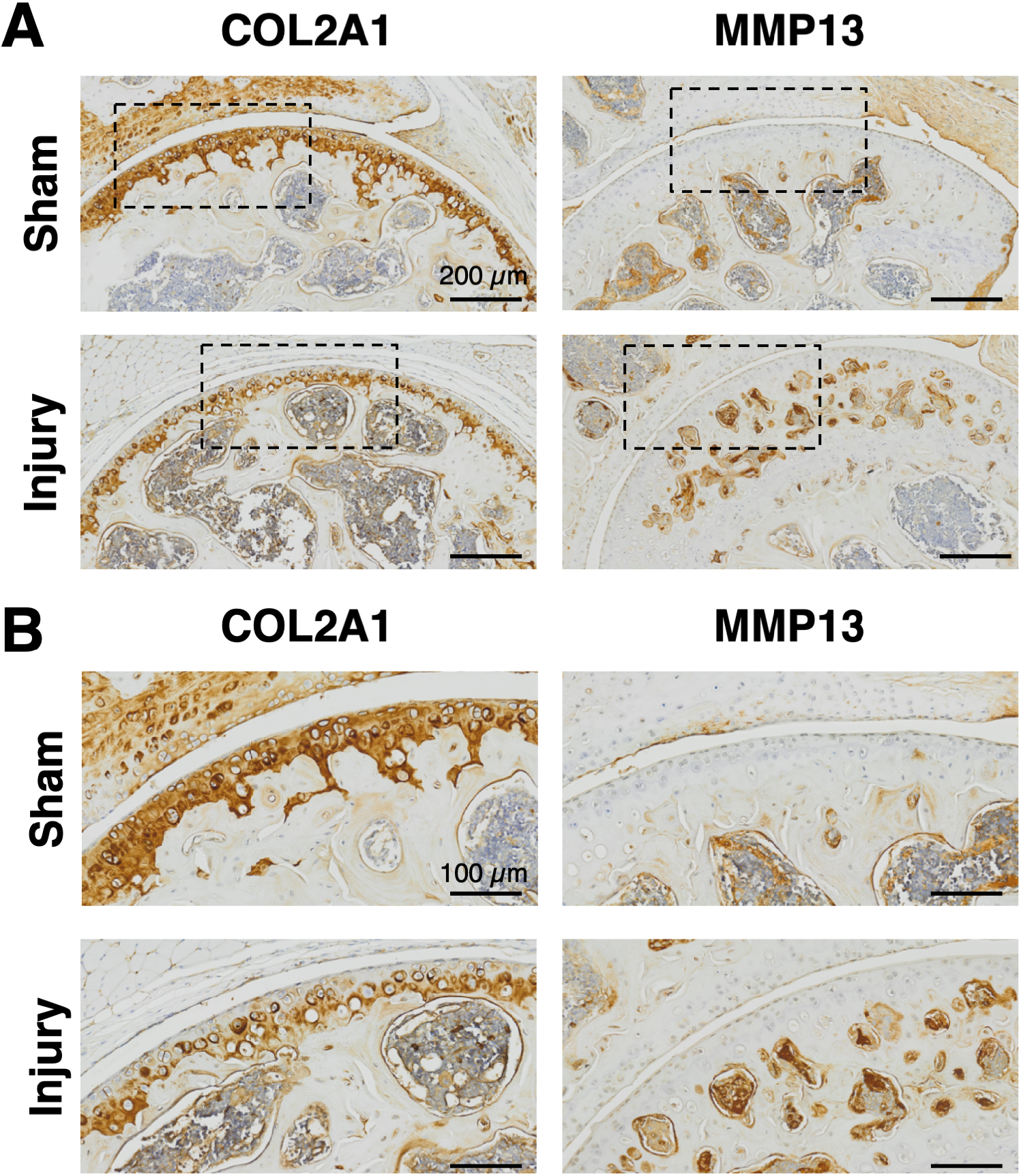
COL2A1 and MMP13 staining reveal loss of type II collagen content in the unmineralized hip cartilage of the mice 20 weeks post-surgery. **(A)** Coronal sections through the hip are shown following IHC for COL2A1 and MMP13 staining. Dark red/brown color indicates positive staining. A substantially decreased COL2A1 staining in the unmineralized hip cartilage of the mice receiving surgery; however, no apparent staining for MMP13 was detected for both sham and surgery groups. (**B**) Increased magnification of the areas indicated by dashed rectangles in (**A**).

MMP13 has been shown to play a catabolic role in OA^19^, and increased staining of MMP13 has been observed in animal models of arthritis^20,21^. IHC staining for MMP13 was performed to visualize changes in femoral head articular cartilage after sham surgery and abductor injury. There was no appreciable staining for MMP13 in the articular cartilage at 20 weeks post-injury in the both sham and surgery groups (**Fig. 3**).

### μCT reveals no significant change in subchondral bone parameters 20 weeks post-injury

To evaluate potential structural changes in subchondral bone, hips from the sham group and the injury group were analyzed by μCT to compare subchondral microtrabecular volumetric parameters 3- and 20-weeks following injury (**Fig. 4**). No changes in cancellous bone fraction (bone volume/total volume, BV/TV, excluding the cortex) were detected in the injured group relative to sham at 20 weeks (Sham: 0.51 ± 0.1; Injury: 0.51 ± 0.1) (**Fig. 4A**). In addition, there were no significant differences in trabecular bone characteristics between two groups. This includes trabecular number (Sham: 6.3/mm ± 0.8; Injury: 6.3/mm ± 0.4) (**Fig. 4B**), trabecular thickness (Sham: 79.2 μm ± 8.1; Injury: 81.8 μm ± 7.3) (**Fig. 4C**), and trabecular spacing (Sham: 151.8 μm ± 23.5; Injury: 147.5 μm ± 10.8) (**Fig. 4D**).

**Figure 4.**
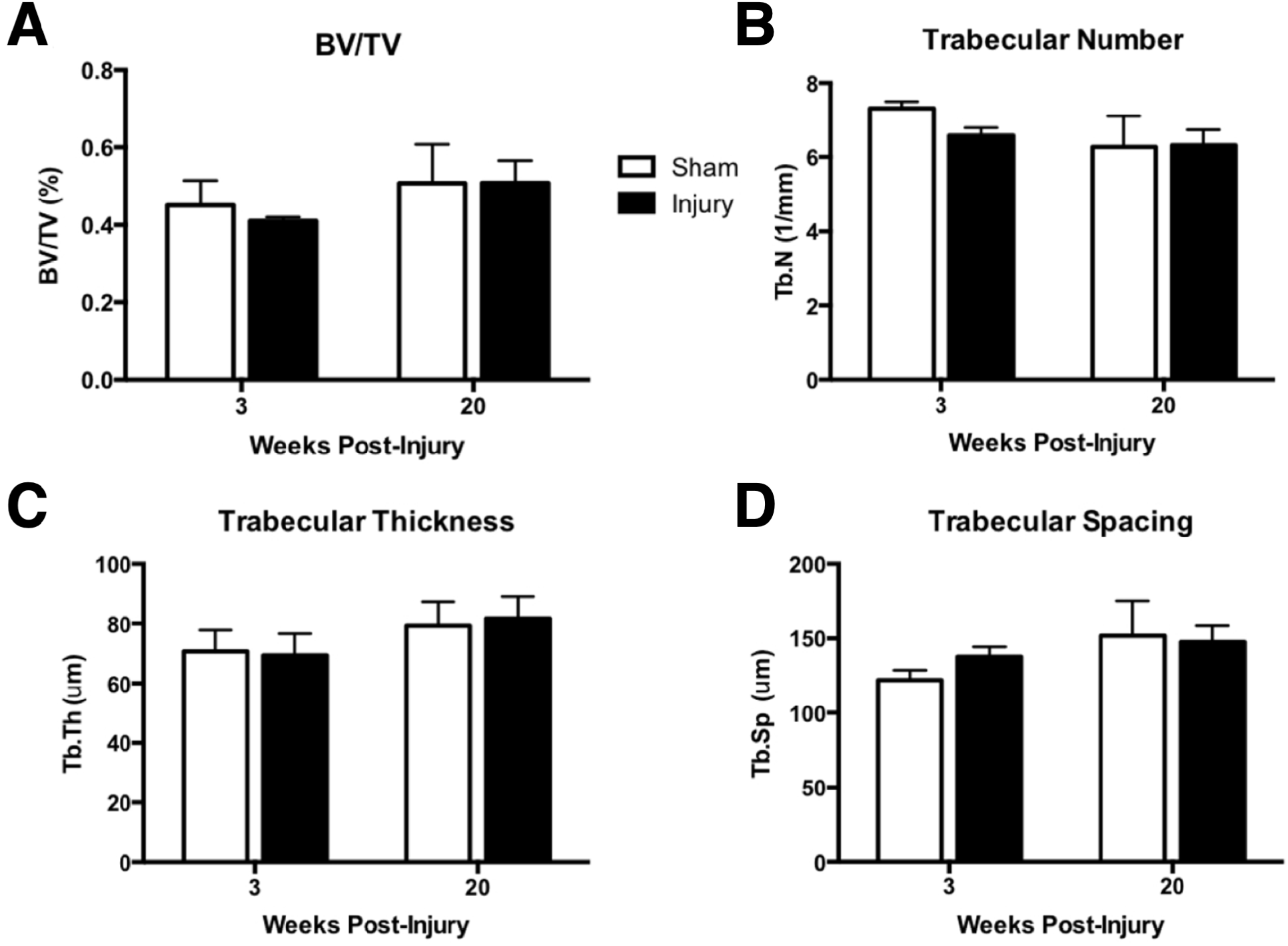
μCT analysis indicates no significant changes in subchondral bone parameters following abductor injury. μCT was used to assess the subchondral bone of the femoral head. (**A**) The ratio of bone volume to total volume (BV/TV), (**B**) Trabecular number, (**C**) trabecular thickness, and (D) trabecular spacing were measured at 3- and 20-weeks post-surgery for bot sham and surgery groups. There are no significant differences in subchondral bone parameters between the two groups in any time point investigated. Two-way repeated measured ANOVA following by Sidak’s multiple comparisons test. Mean ± SD.

## Discussion

With the growth in clinical evidence indicating an association between abductor insufficiency and the development of hip OA, it is essential to develop a reproducible small animal model that allows scientists to study the pathogenesis of this linked disease process^5,6^. To date, however, only a few rodent hip OA models have been established and can be generally categorized into either chemically-induced or surgically-induced OA^16,22^. Intra-articular injection of monosodium Iodoacetate (a chondrocyte glycolytic inhibitor) exhibited rapid hip OA development compared to controls within 14 days in a rat model. However, it is not clear whether such a progressive and destructive OA phenotype is representative of chronic hip OA in humans^22^. In another study, various degrees of hip instability ranging from mild, moderate, severe to femoral head resection were surgically induced in mice at weaning (3-week-old neonatal pups)^16^. Their data suggest that hip instability induced by loss-of-function of soft connective tissue led to morphometric changes in the growing mouse hip. However, understanding how abductor injury may induce OA in the context of a skeletally mature hip (as indicated fusion of triradiate cartilage of the pelvis)^23^, has remained a key knowledge gap to this point. Furthermore, our small animal hip OA model may be used to explore the sequence of pathoanatomical and anabolic/carbolic effects on OA onset and progression.

Animal models of surgical OA require well-established techniques that can be easily reproduced^13^. In this model the third trochanter (3T) serves as a surgical landmark, which can be palpated before making the skin incision. Once the muscular attachments to the 3T have been cut, the femur is followed proximally to the large abductor complex on the greater trochanter (GT). These muscles are easily visualized, and can be cleanly cut without injuring surrounding structures. The surgery avoids dissecting around deep structures; the major potential injury is to the sciatic nerve (SN) running posterior to the proximal femur. In preliminary work, we experimented with performing a capsulotomy to induce an even greater severity of injury.

Through multiple surgeries, we found that the capsule in the mouse may be too small to reliably incise without risking damage to the sciatic nerve running in close approximation to the joint. As such, this group has been excluded from the presented results.

The primary findings in this study were the loss of proteoglycan and COL2A1 staining in uncalcified articular cartilage 20 weeks after injury. Loss of proteoglycan and type II collagen contents are both histologic hallmarks of OA, and the observations made in the injured group after 20 weeks suggest that OA progression is accelerated after abductor injury. Additionally, sham mice had similar OA severity at 3 weeks and 20 weeks post-surgery, suggesting that both sham surgery and aging (i.e. ∼ 6 moths of age) were not sufficient to induce hip OA onset.

Interestingly, no changes in MMP13 IHC staining were observed between the sham and injured groups, suggesting that either MMP13 may not be the main catabolic mediator in our mouse hip OA model or other timepoints may need to be investigated in order to establish temporal expression pattern of MMP13. Indeed, our results are also consistent with the findings from the study of Killian *et al*. where the authors indicated low MMP13 staining in articular cartilage in their titrated model of hip dysplasia. Future studies investigating other cartilage degradation enzymes, such as ADAMTS4, MMP3, MMP9, etc., as well as additional timepoints post-surgery are warranted. Radiographic analysis of subchondral bone remodeling did not reveal significant differences between sham and injured groups at any time point investigated in this study. Nevertheless, long-term study may be required to observe subchondral bone remodeling in our mouse model.

The gluteus medius and minimus, in conjunction with the tensor fascia lata and gluteus maximus, represent the main components of the hip abductor complex. This complex contributes to the “contractile layer” of the hip, and is thought to exert considerable influence on the intraarticular joint space and its associated layers^24^. In a clinical outcome study that evaluated the results of arthroscopic gluteus medius/minimus repairs, a high incidence of concomitant intraarticular hip pathology was noted^7^. Abductor tendon pathology leads to considerable pain and functional impairment. Both open and arthroscopic repair techniques have been proposed and excellent clinical results have been reported for both repair types ^7,25,26^. The causal relationship between periarticular myotendinous pathology and the onset and progression of OA is intriguing but remains understudied. Emerging clinical data for the hip joint support the association between pathology of the extra-articular soft tissue envelope and intra-articular joint disease. Therefore, reestablishment of an effective abductor force coupling mechanism may help to optimize and re-balance loading characteristics and potentially influence the progression of intraarticular degenerative changes that may ultimately result in the need for joint arthroplasty.

The true incidence of abductor tears and/or tendinosis in the general and athletic population remains unknown. In addition, the natural history of untreated tears is unclear. Some studies have observed an association between hip abductor disease and OA ^5,27^. A recent histologic study evaluated the ultrastructure of the abductor tendon complex in 10 patients treated for displaced femoral neck fractures and 10 patients undergoing total hip arthroplasty for OA. All of the patients treated for OA were found to have coexisting tendinosis, with prominent scarring and overall greater degenerative changes than the group treated for traumatic fractures of the femoral neck^6^. While not clearly establishing that tendon pathology contributes to OA, these observations raise important questions regarding the relationship between these two pathologies. It is likely that abductor compromise, through an analogous pathomechanical process, may contribute to the onset and progression of hip OA.

While our present study establishes a reproducible injured-induced mouse hip OA model and offers an important step forward in investigating the role of abductor insufficiency in the development of hip OA, there are limitations that must be considered. First, the mechanisms underlying myotendinous pathology and abductor insufficiency leading to mouse hip OA development remain to be determined. Particularly, to what extent the biomechanical loading on the hip cartilage and mouse behaviors such as gaits and weight bearing are altered following abductor surgery need to be quantified in the future. An additional limitation that must be considered is the clinical relevance of a quadruped model for hip OA, as weight bearing is distributed across four limbs in mice, and thus, gait and biomechanics differ from humans. Despite this limitation, mice and humans share similar joint congruence at the hip.

In conclusion, we have established a novel murine model of surgically-induced hip OA through an injury to the abductor complex around the hip. Furthermore, this model may be utilized as a valuable tool to examine potential biomarkers associated with hip OA progression^28^. We believe that for the broader orthopaedic community of clinicians and researchers, our findings will increase understanding the role of abductors in hip stability and the development of joint pathology.

## Acknowledgements

We would like to thank the Histology, Biochemistry and Molecular Imaging (HBMI) and Biomechanics, Biomechanics and Multimodal Tissue Imaging (BBMTI) cores in the Center for Musculoskeletal Research (CMSR) for technical assistance. We appreciate the gracious generosity of the Goldstein Family, who supported the Goldstein Award which provided funding for this work. We also thank Dr. Victoria Hansen and Gulzada Kulzhanova’s assistance on performing modified Mainkin score grading for hip joint OA severity for the current project.

## Author contributions

AL, CLW, and BG conceptualized the study. MBG, CAO, and JR performed mouse surgeries and μCT measurements. HS, JR, CLW, and members in CLW’s lab performed modified Mainkin score grading for hip joint OA severity. HS performed IHC staining and imaging. All authors wrote, reviewed, and edited the manuscript.

